# A copper switch for inducing CRISPR/Cas9-based transcriptional activation tightly regulates gene expression in *Nicotiana benthamiana*

**DOI:** 10.1101/2021.09.07.459151

**Authors:** Elena Garcia-Perez, Borja Diego-Martin, Alfredo Quijano-Rubio, Elena Moreno-Gimenez, Diego Orzaez, Marta Vazquez-Vilar

## Abstract

CRISPR-based programmable transcriptional activators (PTAs) are used in plants for rewiring gene networks. Better tuning of their activity in a time and dose-dependent manner should allow precise control of gene expression. Here, we report the optimization of a Copper Inducible system called CI-switch for conditional gene activation in *Nicotiana benthamiana*. In the presence of copper, the copper-responsive factor CUP2 undergoes a conformational change and binds a DNA motif named copper-binding site (CBS). In this study, we tested several activation domains fused to CUP2 and found that the non-viral Gal4 domain results in strong activation of a reporter gene equipped with a minimal promoter, offering advantages over previous designs. To connect copper regulation with downstream programable elements, several copper-dependent configurations of the strong dCasEV2.1 PTA were assayed, aiming at maximizing activation range, while minimizing undesired background expression. The best configuration involved a dual copper regulation of the two protein components of the PTA, namely dCas9:EDLL and MS2:VPR, and a constitutive RNA pol III-driven expression of the third component, a guide RNA with anchoring sites for the MS2 RNA-binding domain. With these optimizations in place, the CI/dCasEV2.1 system resulted in copper-dependent activation rates of 2,600-fold for the endogenous *N. benthamiana* DFR gene, with negligible expression in the absence of the trigger. The tight regulation of copper over CI/dCasEV2.1 makes this system ideal for the conditional production of plant-derived metabolites and recombinant proteins in the field.

## BACKGROUND

Synthetic biology offers the opportunity to overcome intrinsic limitations of biological systems for biotechnology applications by precisely rewiring genetic circuits. Standardization and modularization of DNA elements lead to the creation of synthetic circuits that can orchestrate the endogenous circuitry in the plant (Liu & Stewart, 2015; Nandagopal & Elowitz, 2011). The implementation of effective synthetic circuits relies on universal devices that incorporate properly characterized genetic elements. Transcriptional regulation based on CRISPR/Cas9 technology has become a popular tool for controlling gene expression in eukaryotes (Konermann *et al*., 2015; Tanenbaum *et al*., 2014; Zhang *et al*., 2015). CRISPR-based genetic circuits provide a precise solution for connecting synthetic devices with endogenous networks and pathways. The binding-ability customization of the nuclease-inactivated Cas9 protein (dCas9) by tailoring guide RNAs allows targeting DNA sequences with high precision (Zhang, 2019). The customizable dCas9 can be linked to transcriptional activation domains leading to the so-called programmable transcriptional activators (PTAs). The use of these synthetic PTAs leads to advanced control over gene expression across diverse plant systems (Pickar-Oliver & Gersbach, 2019).

We previously developed a potent PTA named dCasEV2.1 (Selma *et al*., 2019). This PTA comprises two transcriptional units (TUs) for expressing two protein components and a third TU for the guide RNAs expression cassette. The first protein component is the dCas9 fused to the EDLL transcriptional activation domain (Tiwari *et al*., 2012). The second component is a fusion protein made of the MS2 phage coat protein and a VPR activation domain. VPR is a synthetic cluster of viral activators that further recruits the cell machinery for initiating transcription (Chavez *et al*., 2015). The third component of the dCasEV2.1 system is the guide RNA (gRNA) that targets the dCas9 to a specific position in the DNA and includes an aptamer in its scaffold for the binding of MS2 (Kunii *et al*., 2018). Together, EDLL and VPR, have a synergic effect on the activation of the targeted gene leading to high expression levels (Selma *et al*., 2019).

Metabolic engineering and molecular farming usually aim at maximizing the expression levels of genes of interest. However, this maximization is often accompanied by serious detrimental effects such as toxicity or developmental defects, which can be mitigated by engineering tight control systems (Bernabé-Orts *et al*., 2020). The implementation of genetic switches actuating over PTAs would permit the creation of inducible activation circuits thus resulting in a fine-tuned regulation of genetic networks (Mandegar *et al*., 2016). Inducers act as stimuli for sensor proteins. These sensors convert a particular stimulus into a transcriptional signal that leads to the activation of a downstream gene. Depending on the type of stimulus, different inducible PTA systems can be established that allow the expression of the dCas9 activator under a specific condition. Traditional inducible systems to control transcription in plants are based on stimuli like chemicals, light, temperature, hormones, stress and wound (*Tang et al*., 2004). Most of these inducers can trigger deleterious pleiotropic effects in the plant chassis and are complicated to apply and monitor in large crop fields (Corrado & Karali, 2009). However, among chemical inducers, micronutrients such as copper represent an innocuous and inexpensive group.

Copper is commonly applied to crops as copper sulfate (CuSO_4_) not only like an essential micronutrient but also like fungicide. Copper sulfate can be easily taken up by plants, it is low-cost, easy to apply and it is already registered for field use as a fungicide (Kumar *et al*., 2021). Accordingly, copper represents a potentially advantageous inducer for programmable transcriptional activation systems. Mett *et al*. (1993) developed a gene expression system inducible by copper based on the copper-metallothionein regulatory system from yeast. This regulatory system stands on a Cu^2+^-regulated transcription factor from *Saccharomyces cerevisiae* (Buchman *et al*., 1989). In the presence of the trigger, the yeast copper responsive factor CUP2 (also known as ACE1) undergoes a conformational change that enables it to specifically bind a DNA operator sequence known as Metal Responsive Element (MRE) or Copper Binding Site (CBS). The binding of CUP2 to a CBS adjoining a minimal promoter leads to the transcriptional activation of the downstream gene. The copper-inducible system initially described by Mett *et al*. (1993) was later optimized by (Saijo & Nagasawa, 2014). This optimization consisted of the concatenation of four repetitions of the CBS for recruiting more molecules of CUP2. Also, a minimum sequence of the 35S promoter was reported to reduce the leaky expression of the targeted gene. Finally, the viral transcription activator VP16 was fused to CUP2 to boost the activation capacity of CUP2. This renewed system worked *in planta* and overcame the copper-inducible gene expression results reported earlier.

Here we show an optimized Copper-Inducible gene expression system named CI-switch, which incorporates the Gal4 yeast activator domain fused to CUP2. The CI-switch is free of viral elements and shows considerably reduced background expression in the absence of copper. Moreover, we show the implementation of a small copper-triggered customizable activation cascade that connects the CI-switch to the expression of dCas9EV2.1, with the subsequent activation of either reporters or endogenous genes as downstream elements in *Nicotiana benthamiana*.

## RESULTS

### Optimization of a copper responsive genetic switch for efficient reporter gene activation in *N. benthamiana*

To develop an agronomically compatible system for switching on transcription, we explored the optimization of a copper-dependent system in *N. benthamiana*. Four repeats (4x) of the Copper Binding Site (CBS) operator were assembled upstream a minimal promoter of the *Solanum lycopersicum* NADPH-dependent dihydroflavonol reductase (DFR) gene for assessing the transcriptional activation of Firefly luciferase (FLuc) reporter gene (Figure 1A). This minimal promoter spans the 155 bp upstream of the TSS of the DFR promoter, which is characterized by a low and non-variable expression (Selma *et al*., 2019). This promoter is hereafter referred as CBS:minDFR. The copper responsive transcription factor CUP2 was expressed under the constitutive Nopaline synthase promoter (pNOS) and fused to different transcriptional activation domains. We compared the performance of the yeast Gal4 activation domain with the activation provided by the viral-based domains VP16, VPR (VP64-p65-Rta), and TV (6x TAL, 4x VP64), commonly used as strong activation elements in plants (Casas-Mollano *et al*., 2020). Copper sulfate 5 mM was sprayed onto the leaves to induce the transiently expressed system. Luminescence assays showed that the basal expression of the reporter gene in the absence of copper resulted in negligible levels in all tested activation fusions (Figure 1B). The relative transcriptional activities (RTAs) conferred by the CBS:minDFR promoter after copper-mediated activation ranged from 4.5±1.5 to 13.5±6.2, as compared with the expression levels conferred by a pNOS promoter in the same assay conditions (Vazquez-Vilar *et al*., 2017). VPR was reported as the strongest activator, but also as the most variable one. The RTAs obtained with Gal4, VP16, and TV in the presence of copper were not statistically significant between each other. Therefore, Gal4 resulted as a suitable alternative to viral domains in the copper-responsive genetic switch. We named this novel combination of the copper-responsive factor CUP2:Gal4 and the 4xCBS operator upstream the minimal DFR promoter as CI-switch.

**Figure 1.**
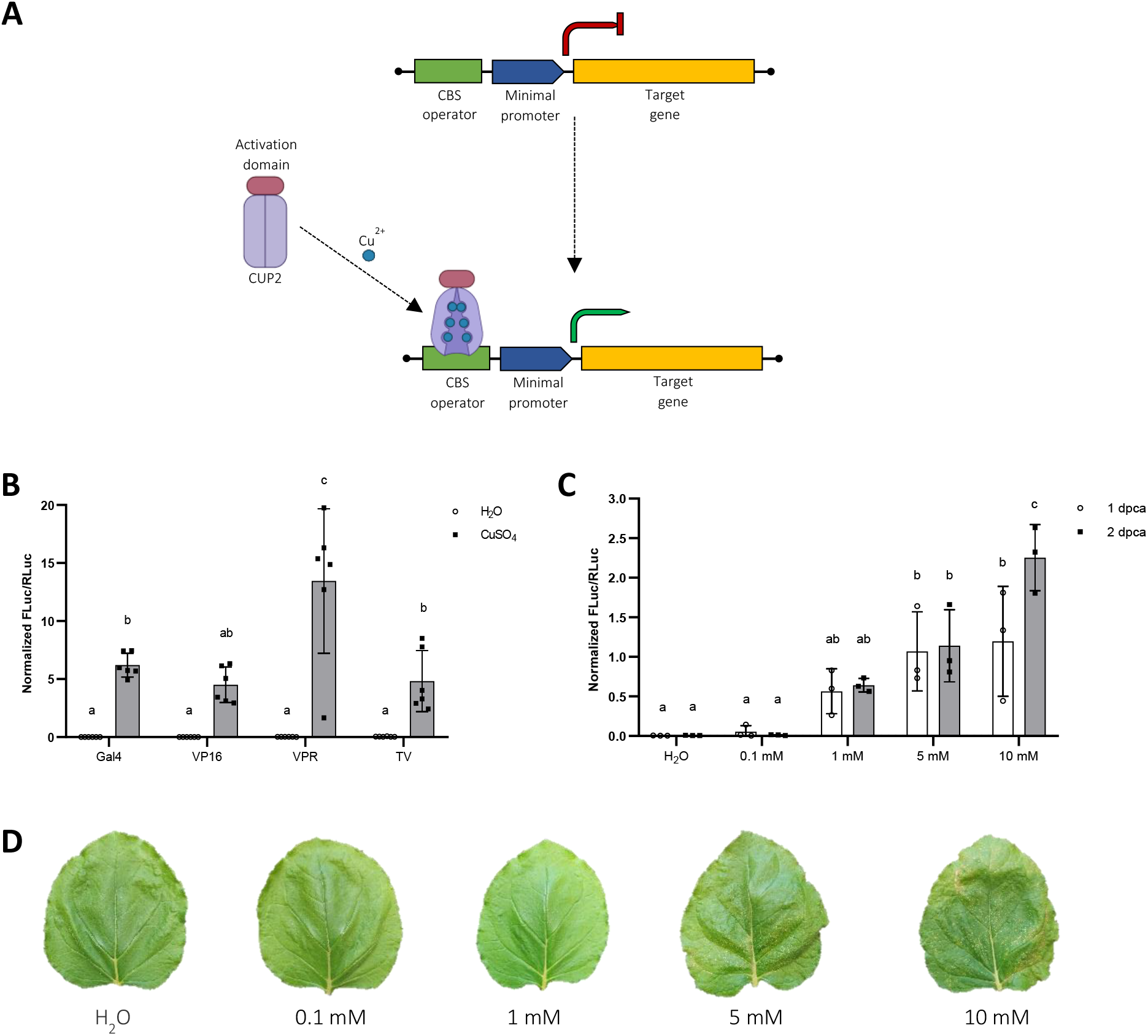
Copper-inducible reporter gene activation in transient expression in *Nicotiana benthamiana*. **A)** Schematic representation of the copper-inducible gene expression system. The CUP2 fused to an activation domain such as Gal4 changes its conformation upon copper binding and binds a Copper Binding Site (CBS) operator allowing transcription of the downstream gene (green arrow). **B)** Copper-mediated activation of Firefly Luciferase (FLuc) reporter gene by different activation domains (Gal4, VP16, VPR, and TV) fused to CUP2 after 5 mM copper sulfate application. **C)** Copper-mediated activation of FLuc by different concentrations of copper sulfate. The construct CUP2:Gal4 was used as the copper-dependent transcriptional factor. **D)** Pictures of leaves 5 days after treatment with different concentrations of copper sulfate. For B) and C), activation is expressed as Normalized FLuc/RLuc ratios of *N. benthamiana* leaves transiently expressing the genetic constructs. Error bars indicate SD (n≥3). Statistical analyses were performed using one-way ANOVA (Tukey’s multiple comparisons test, P-Value ≤ 0.05). Variables within the same statistical groups are marked with the same letters. Figure includes images from Biorender (biorender.com).

Next, we optimized the copper concentrations used for the CI-switch. For this, the sprayed CuSO_4_ volume per leaf was set to 5 ml and different concentrations of copper sulfate (CuSO_4_) were tested. Also, the FLuc activity used as standard reference was analyzed at two time points, one- and two days post-copper-induction, to determine whether the system could result in higher activations rates at times longer than 24 h. Figure 1C shows the dose-dependent response in the system when applying different concentrations of CuSO_4_. The basal expression of the system without copper was virtually null, thus leakiness of the CI-switch was not observed. The system was not activated with 0.1 mM copper, but higher levels of Fluc expression were induced with increasing CuSO_4_ concentration. Maximum RTA values of 2.3±0.4 were obtained with 10 mM CuSO_4._ However, at this concentration necrosis symptoms in the leaf tissue were noticeable (figure 1D). To avoid any potential phytotoxic effect, we selected 5 mM for subsequent experiments. Differences in FLuc activity between one- and two-days post-copper application (dpca) were not statistically significant at any of the assessed concentrations except for the detrimental 10 mM. Therefore, the system does not require more than one day to reach its maximum functionality.

### Optimization of dCasEV2.1 copper-mediated regulation in *N. benthamiana*

To assess the modulation that copper can offer in the regulation of transcriptional gene circuits, we connected the copper switch to the three-component dCasEV2.1 PTA, and tested the performance of the resulting genetic device using a transient luciferase assay in *N. benthamiana*. The CI/dCasEV2.1 tool comprised the CUP2:Gal4 TU, the 3X TUs dCasEV2.1 complex (dCas9:EDLL, MS2:VPR and the adapted gRNA) (figure 2A, B), and a FLuc reporter TU that contains the *S. lycopersicum* DFR promoter targeted by the gRNA. We employed two different copper-regulated synthetic promoters, namely CBS:min35S (carrying the 4xCBS box next to a minimal 35S promoter), and the previously tested CBS:minDFR. In our first optimization steps, the CUP2:Gal4 and the gRNA TUs were constitutively expressed (driven by pNOS and *Arabidopsis thaliana* U626 respectively), and the remaining elements were assayed under different copper inducible configurations. In the Cu/Cs configuration, only dCas9:EDLL TU was under copper regulation, whereas MS2:VPR TU was constitutively expressed (figure 2B). In the Cu/Cu configuration both TUs were put under copper regulated promoters. Both Cu/Cs and Cu/Cu configurations were assayed using the two synthetic promoters described above. As can be observed in figure 2C, when the minDFR element was present, the single Cu configuration (Cu/Cs) showed moderate background levels (0.16±0.04), which were almost completely abolished when dual copper regulation (Cu/Cu) was implemented (0.01±0.05). Furthermore, dual regulation offered higher activation rates of FLuc reporter than single regulation, reaching average RTAs of 1,43±1.00 (figure 2C).

**Figure 2.**
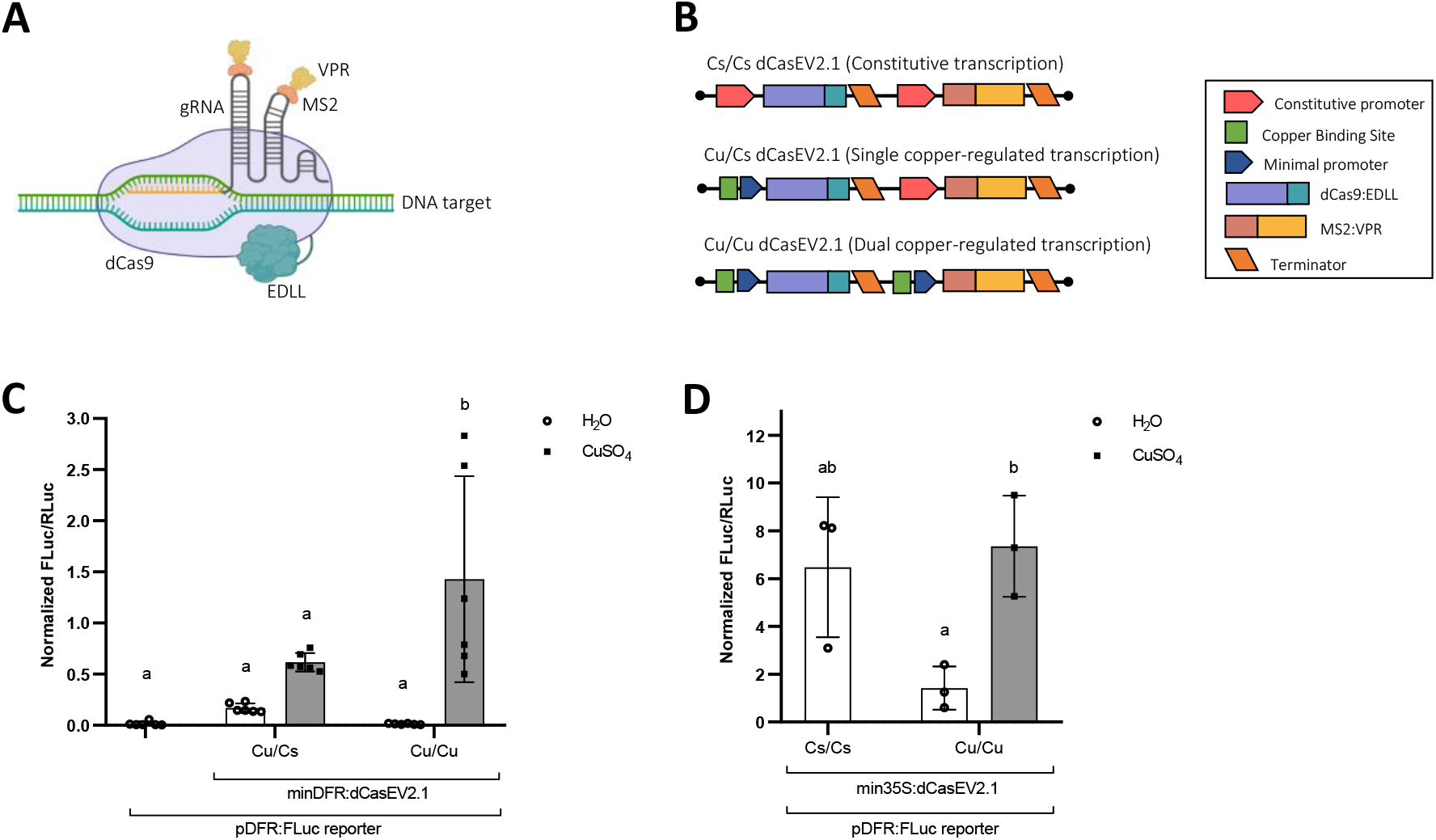
Copper-inducible reporter gene activation by dCasEV2.1 in *N. benthamiana*. **A)** Schematic representation of dCasEV2.1 components. **B)** Genetic constructs for dual constitutive (Cs/Cs) transcription (35S promoter for both dCas9:EDLL and MS2:VPR), single copper-regulated (Cu/Cs) transcription (CBS:minimal DFR promoter for dCas9:EDLL and 35S promoter for MS2:VPR), and dual copper-regulated (Cu/Cu) transcription (CBS:minimal DFR promoter for both dCas9:EDLL and MS2:VPR). **C)** Efficiency of single vs. dual copper-regulated transcription of dCasEV2.1 under a minimal DFR promoter (minDFR) to activate a pDFR:FLuc reporter. **D)** Efficiency of dual copper-regulated transcription of dCasEV2.1 under a minimal 35S (min35S) promoter to activate a pDFR:FLuc reporter. Activation is expressed as Normalized FLuc/RLuc ratios of *N. benthamiana* leaves transiently expressing the genetic constructs. All FLuc/RLuc ratios were normalized using the FLuc/RLuc ratios of *N. benthamiana* leaves expressing a constitutive pNOS:Fluc reporter. Error bars indicate SD (n≥3). Statistical analyses were performed using one-way ANOVA (Tukey’s multiple comparisons test, P-Value ≤ 0.05). Variables within the same statistical groups are marked with the same letters. Figure includes images from Biorender (biorender.com).

The same analysis was performed using the CBS:min35S synthetic promoter (Saijo & Nagasawa, 2014). As shown in figure 2D, the RTAs achieved by using min35S-containing constructs were consistently higher than those obtained with minDFR and even slightly higher than with the constitutive version, Cs/Cs (figure 2D). However, constructs employing min35S resulted in higher basal expression and there is no statistically significant difference in the activation provided by the constitutive dCasEV2.1 and the Cu/Cu configuration when copper is not present.

### Effect of gRNA regulation on copper-inducible dCasEV2.1 activation

Once the regulation of the dCasEV2.1 protein components was optimized, we aimed to further explore the control of the gRNA expression with copper and ethanol. The choice of a second inducer for gRNA expression aims at enabling the activation of different set of genes under different stimuli. CRISPR/Cas9 guide RNAs are usually transcribed under a RNA polymerase III (pol-III) promoter, like the U626 (Cong *et al*., 2013; Mali *et al*., 2013). However, most minimal promoters coupled to inducible operators rely on polymerase II (pol-II) transcription (Knapp *et al*., 2019). As pol-II transcripts are altered with a 5’ cap and poly-A tail and exported from the nucleus, the efficiency of a gRNA expressed under pol-II would be highly compromised. Thus, we flanked the sequence of the gRNAs by the *A. thaliana* tRNA^Gly^ sequence (Xie *et al*., 2015) to ensure the post-transcriptional processing of the 5’ and 3’ ends generated after pol-II action. To assemble pol-II gRNA expression cassettes using the type IIS DNA assembly system GoldenBraid, we followed the strategy previously described by (Vazquez-Vilar *et al*., 2016) for pol-III gRNAs. In this strategy, the gRNA unit is assembled in a BsmBI restriction-ligation reaction as a Level 0 part incorporating the protospacer sequence as a primer dimer with unique 4-bp overhangs and the 5’ and 3’ sequences flanking the protospacer are provided from another vector. This vector carries (i) the tRNA^Gly^ sequence and (ii) the scaffold-tRNA^Gly^ sequence, both flanked by BsmBI restriction sites and unique overhangs for their assembly upstream and downstream the protospacer sequence, respectively. In the next step the gRNA unit is assembled between a Pol-II promoter and a transcriptional terminator in a BsaI restriction-ligation reaction (figure 3A). A new webtool has been developed during this work to design and assemble tailored PolII polycistronic gRNA constructs for gene regulation with (d)Cas9 (https://gbcloning.upv.es/tools/cas9multiplexing_inducible_regulation/).

**Figure 3.**
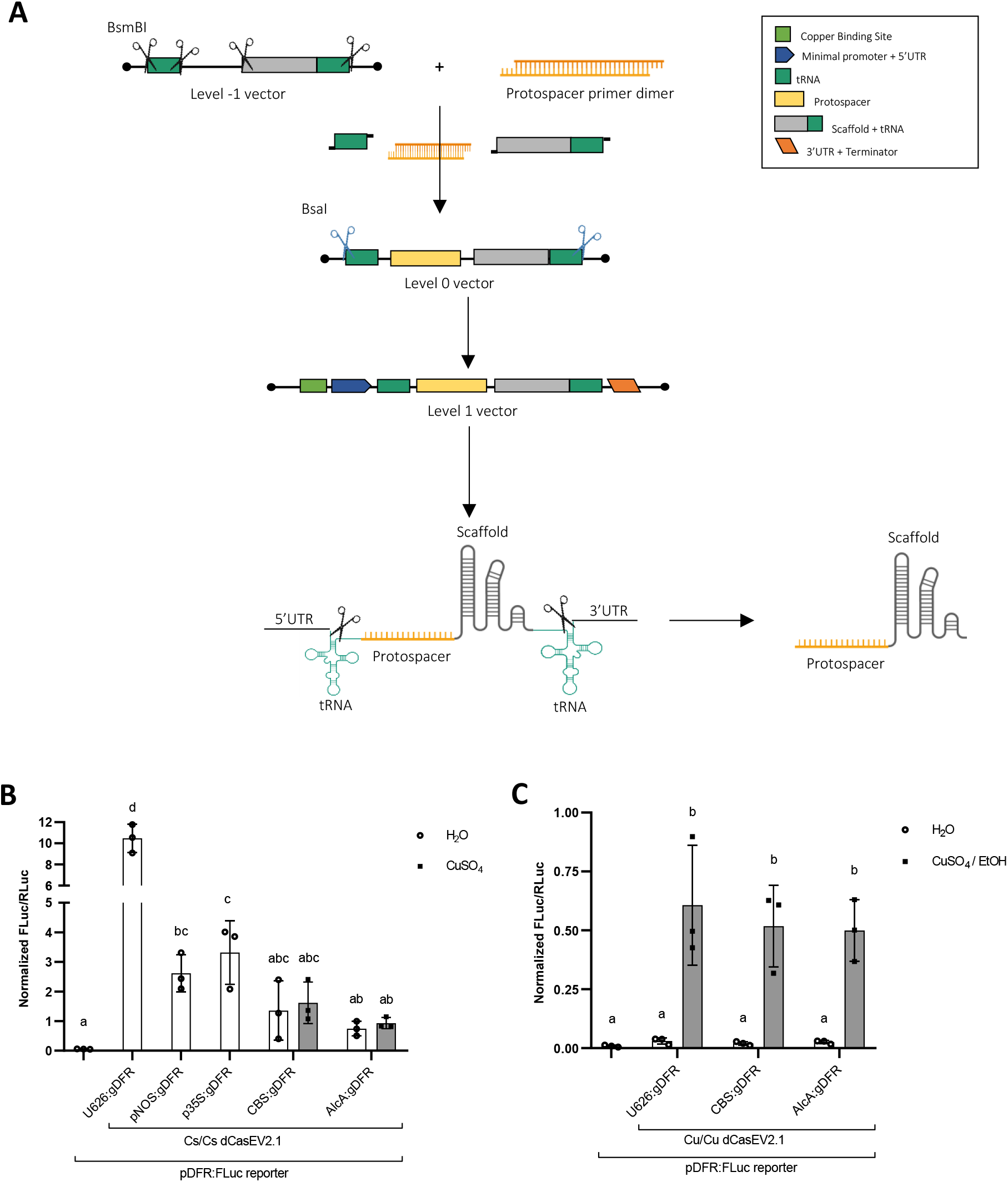
Reporter gene activation by chemically inducible pol-II expressed guide RNAs in combination with dCasEV2.1. **A)** Schematic representation of cloning strategy and post-transcriptional processing of guide RNA (gRNA). **B)** Reporter gene activation with constitutive dCasEV2.1 and differently-expressed gRNAs. The pDFR:FLuc reporter was activated by a constitutive dCasEV2.1 under two 35S promoter (Cs/Cs dCasEV2.1) combined with differently expressed gRNAs targeting the DFR promoter of the FLuc reporter. U626 is a constitutive promoter for polymerase III, and pNOS and p35S are constitutive promoters for polymerase II. CBS is the operator for copper-induction and AlcA is the operator for ethanol-induction. Both CBS and AlcA are assembled next to a minimal DFR promoter. **C)** Triple induction of dCasEV2.1 components to activate gene expression. The pDFR:FLuc reporter was activated by a double-induced dCasEV2.1 using a minDFR promoter (Cu/Cu dCasEV2.1) in combination with differently expressed gRNAs targeting the DFR promoter of the FLuc reporter. Activation is expressed as Normalized FLuc/RLuc ratios obtained for the *N. benthamiana* leaves transiently expressing the genetic constructs. All FLuc/RLuc ratios were normalized using the FLuc/RLuc ratios of *N. benthamiana* leaves expressing a constitutive pNOS:Fluc reporter. Error bars indicate SD (n=3). Statistical analyses were performed using one-way ANOVA (Tukey’s multiple comparisons test, P-Value ≤ 0.05). Variables within the same statistical groups are marked with the same letters. Figure includes images from Biorender (biorender.com).

To assess the feasibility of expressing gRNAs by the pol-II, two constitutive promoters for the pol-II, pNOS and p35S, were tested upstream the tRNA-protospacer-scaffold-tRNA. To evaluate the regulation capability over the pol-II-synthesized gRNAs, the synthetic CBS:minDFR promoter was used for copper-regulation. In parallel ethanol regulation was attempted as well, using AlcR as the ethanol-responsive transcriptional factor that binds to AlcA operator in presence of ethanol (Roslan *et al*., 2001). All the constructs for Pol-II-synthesized gRNAs contained as, in previous experiments, a protospacer targeting a DFR promoter upstream of the reporter FLuc gene. The different gRNA constructs were co-infiltrated with the construct for the constitutive expression of dCasEV2.1, the reporter construct, and the responsive-transcription factor in the plant. When pNOS and p35S were used for gRNA synthesis, activation of FLuc by dCasEV2.1 reached RTAs of 2.6±0.6 and 3.1±1.1 respectively (figure 3B). However, this activation is significantly lower (approximately 3.5-fold) than the activation obtained when the gRNA was synthesized under the pol-III promoter U626. Unfortunately, when gRNAs were controlled by CBS or AlcA operators, FLuc reporter activity reached similar rates both in presence and absence of copper/ethanol.

Our results indicated that the chemical induction of the gRNA component of dCasEV2.1 alone was not sufficient to regulate downstream gene expression. A possible explanation is that gRNA differential expression, if leaky, may be masked by the high and constitutive expression of the other two components in the system. For this reason, we explored the performance of a triple-controlled system combining the double copper-inducible dCasEV2.1 with either a CBS:minDFR:gRNA:Tnos or an AlcA:minDFR:gRNA:Tnos. In all combinations assayed, reporter activation in presence of the inducer was observed. However, as expected, transient-expression assays showed that triple regulation (Cu/Cu/EtOH or Cu/Cu/Cu) gave similar results to those observed with the dCasEV2.1 Cu/Cu conformation, as no differences were found when gRNAs were expressed under the constitutive U626 promoter or the copper- or ethanol-inducible promoters (figure 3C). Therefore, we concluded that the most efficient way to regulate the dCasEV2.1 system is via the regulation of its two protein components dCas9:EDLL and MS2:VPR.

### Copper-regulated dCasEV2.1-mediated activation of endogenous genes

Using dCasEV2.1 as an intermediate operator to the copper-responsive factor offers the possibility of transferring copper regulation to any endogenous gene of interest in the plant (figure 4A) only by reprogramming the gRNA component of the dCasEV2.1 complex. To show this, we programmed dCasEV2.1 to target the *N. benthamiana* DFR (NbDFR) gene. This gene is involved in the anthocyanin biosynthesis pathway and its basal expression is very low (Selma *et al*., 2019). A dCasEV2.1 construct in Cu/Cu conformation (CBS4:minDFR:dCas9:EDLL:tNOS-CBS4:minDFR:MS2:VPR:tNOS) was targeted with a pol-III expressed multiplexing gRNA to three positions of the NbDFR promoter (Selma *et al*., 2019). A gRNA targeting the *S. lycopersicum* MTB gene was included as a negative control of the activation. The constitutively expressed dCasEV2.1 was used as a positive control for maximum activation levels. As can be observed in the figure 4B, the spraying of leaves with 5mM copper solution resulted in a remarkable 2600-fold activation of NbDFR mRNA levels. The expression of NbDFR was negligible in absence of copper inducer, an indication of the highly reduced background levels achieved with the optimization of the copper-responsive genetic tool developed in this work.

**Figure 4.**
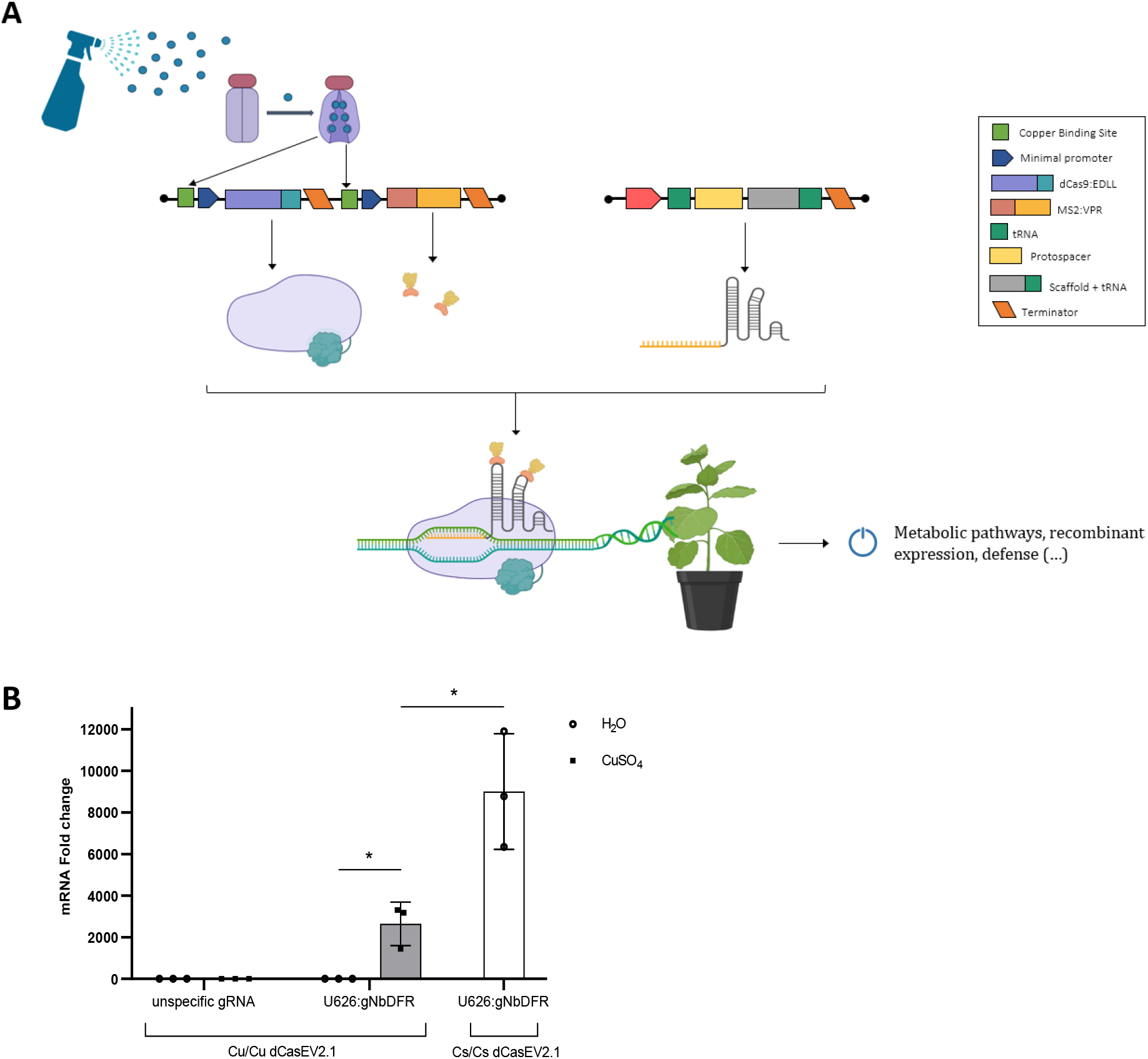
Application of copper-inducible dCasEV2.1 for activating endogenous genes in *N. benthamiana*. **A)** Schematic representation of the strategy of copper-mediated dCasEV2.1 induction for targeted gene activation *in planta*. **B)** Activation of the *N. benthamiana* DFR (NbDFR) gene by the dual copper-regulated dCasEV2.1 under minDFR promoter (Cu/Cu dCasEV2.1) and the constitutive dCasEV2.1 under 35S promoter (Cs/Cs dCasEV2.1). NbDFR transcripts were quantified by RT-qPCR at 5 days post-infiltration of the dually copper-regulated dCasEV2.1 in combination with a constitutive guide RNA targeting either NbDFR (gNbDFR) or the *S. lycopersicum* MTB gene (unspecific gRNA) and a constitutive CUP2:Gal4. Leaves were treated either with 5 mM copper sulfate or water at 3 days post infiltration. Error bars indicate SD (n=3). The asterisks represent a significant difference between treatments. Statistical analysis was performed using unpaired two-tailed t-test for two samples comparisons (P-value ≤ 0.05). Figure includes images from Biorender (biorender.com).

## DISCUSSION

CRISPR-based PTAs are becoming extensively used for endogenous genes activation in plants due to their high specificity and their effectiveness in boosting transcriptional activation (Konermann *et al*., 2015; Tanenbaum *et al*., 2014; Y. Zhang *et al*., 2015). The exploitation of their full potential as tools for the activation on-demand of metabolic and signalling networks relies on their tight regulation. In plants, several chemical inducible systems have been used for controlled gene expression. These systems are triggered by different compounds including antibiotics, ethanol, steroids, or copper (Andres *et al*., 2019). The main advantages of using copper as an inductor for plants are that copper sulfate is cheap and represents a friendly substance in agriculture. Thus, we optimized a copper-inducible system named CI-switch to regulate the programmable transcriptional activator dCasEV2.1 (Selma *et al*., 2019).

Since the first report on the use of the Cu^2+^-regulated transcription factor CUP2 for gene activation in plants (Mett *et al*., 1993), the system has been adapted for different applications. However, most of these applications were not effective enough because of poor induction (Boetti *et al*., 1999; Granger & Cyr, 2000; Mohamed *et al*., 2001), variability between plants, and the leakiness (Granger & Cyr, 2001). (Saijo & Nagasawa, 2014)) boosted the initial system by fusing the viral activation domain VP16 to CUP2 and by incorporating 4 repetitions of the Copper Binding Site (CBS) operator next to a minimal 35S promoter upstream the gene of interest. With this improved system, they reported a high transcription activation of GFP in transgenic lines as well as the induction of flowering in *Arabidopsis thaliana*. In this work, we tested different activation domains in combination with CUP2 to explore a higher activation capacity of the system. Contrary to Saijo & Nagasawa (2014), in our hands, the use of VP16 did not confer a significant improvement to the system. Most interesting, Gal4 fusion resulted in a potent activator. According to current regulations (Turnbull *et al*., 2021), the application of the eukaryotic Gal4 domain at the open field would be more straightforward than the application of the viral VP16. Ultimately, the goal of this work is to provide transcriptional switches which can be realistically deployed in the field, and therefore we decided to continue our characterization with Gal4. We are aware that, despite the choice of Gal4, the system described in this work is not completely free of viral domains, as dCasEV2.1 contains the VPR domain, a tandem of activation domains containing viral sequences (VP64-p65-Rta). While VPR showed the highest activation rates and therefore was the domain of choice for building the dCasEV2.1 tool, other non-viral domains were successfully assayed during dCasEV2.1 optimization which are available in the GoldenBraid collection and the Addgene repository (Selma *et al*., 2019). Although for the optimization experiments described here we chose the use of dCasEV2.1, the alternative non-viral domains are likely to yield similar responses although probably with smaller activation rates. Recently, Pan *et al*. reported that the fusion of two copies of the transcription activator-like effector (TALE) TAL Activation Domain (TAD) to MS2 resulted in strong activation (Pan *et al*., 2021). An alternative configuration for our dCasEV2.1 incorporating TAD may be convenient for developing a free-viral tool to combine with our CI-switch.

When connecting an inducible switch to a potent transcriptional activator, such as dCasEV2.1, it is important to consider that the signal is highly amplified and a minimal leakiness in the inducible system can result in a non-desired activation of the gene of interest in the absence of inductor. Therefore, for the novel application of copper to regulate a PTA, we explored single and dual regulation of both transcriptional units that make up dCasEV2.1. We also investigated two different minimal promoters, DFR, characterized for its low basal expression, and 35S, used in previous studies for copper regulation. The results obtained in this work showed that dual regulation presented lower leakiness and resulted in a higher activation than single regulation. In the case of dual regulation, 110- and 5.18-fold increments of reporter activity were obtained with the minimal DFR and minimal 35S, respectively. To overturn leakiness in the system, minimal DFR resulted as the most suitable promoter to use since the basal expression is practically null. However, if the goal of the activation is reaching a higher expression rate, a minimal 35S promoter would be more promising despite the leakiness. For supplying an extra regulation point to dCasEV2.1 activation, we attempted regulation of the guide RNA. For this regulation, we flanked the gRNA of interest by two tRNA^Gly^ sequences and we tested different pol-II promoters. However, this extra regulation did not lead to efficient control over the activation of the reporter gene. These results are consistent with those reported by Knapp et al. (Knapp *et al*., 2019) showing that tRNAs can generate functional gRNAs that are constitutively and promoter-independently expressed. Knapp *et al*. (2019) designed a tRNA^Pro^ scaffold (Δ, ΔC55A, and ΔC55G) with minimized promoter strength without affecting the processing activity of the tRNA. An interesting addition to our work would be to implement the usage of these tailored tRNAs.

The strict control of an inducible gene expression system is essential when studying signaling networks or when producing recombinant proteins with toxic effects in plants. Thus, our optimization steps of a copper-inducible PTA aimed to get maximum expression levels while keeping low basal expression. Altogether, our results showed that the best configuration for the Copper-Inducible switch consists of a constitutive CUP2:Gal4 transcription factor able to bind and activate the expression of the two protein components of the dCasEV2.1 from a CBS operator and minDFR promoter in the presence of 5 mM copper sulfate. The gRNA targeting the PTA to the gene of interest can be expressed constitutively from a pol III promoter. When using this system for *N. benthamiana* DFR gene activation we observed no leakiness and a 2600-fold transcriptional activation after copper application, thus confirming the robustness of the tool. The stable transformation of the copper-inducible PTA developed in this work along with user-designed gRNAs would result in a highly specific spatial/temporal expression of the genes of interest, either native or transgenes, in the plant. Furthermore, the application possibilities of the system may be expanded with a regulated expression of the gRNAs. This could be achieved either by the incorporation of tRNAs with minimized promoter strength, as discussed previously, or with tissue-specific promoters. The feasibility of regulating gRNA synthesis would open the possibility of creating AND-logic gates to tightly control endogenous gene activation on-demand.

## METHODS

### Cloning and assembly of the GoldenBraid constructs

The constructs used in this work were assembled using the GoldenBraid (GB) cloning system (Sarrion-Perdigones et al., 2013; Vazquez-Vilar et al., 2017). This system is based on subsequent Type IIS-restriction-ligation reactions to clone the desired TUs or combinations of TUs in destination vectors. Shortly, in the GoldenBraid hierarchy, basic DNA parts such as promoters, terminators, and coding sequences of genes of interest are considered as level 0 DNA parts. These level 0 parts are cloned in the pUPD2 plasmid via a BsmBI-mediated restriction-ligation reaction. Transcriptional units represent level 1 and they are assembled via a BsaI-mediated restriction-ligation reaction using a pDGB3 α as destination vector. The transcriptional units are combined to create complex genetic modules in level >1, by using BsaI- and BsmBI-mediated assembly reactions into pDGB3 α and pDGB3 Ω destination vectors. All the constructs used in this work are listed in Supplementary Table S1, as well as their identification numbers to track their sequence and their assembly history at the GoldenBraid repository (https://gbcloning.upv.es/).

### Agroinfiltration experiments

Transient expression of the genetic constructs was performed in wild-type *Nicotiana benthamiana* leaves. *Agrobacterium tumefaciens* strain C58 harboring the expression vectors were grown in LB medium supplemented with 50 μg/ml kanamycin or spectinomycin and 50 μg/ml rifampicin for 16 hours at 28°C/250 rpm. Subsequently, the bacteria were pelleted, transferred to infiltration buffer (10 mM MES, 10 mM MgCl_2_, and 200 μM acetosyringone, pH 5.6), and incubated for 2h at room temperature in a rolling mixer. After this incubation step, the OD_600_ of the cultures was adjusted to 0.1. For plant co-expression of more than one GB element, the different *Agrobacterium* cultures were mixed and co-infiltrated in three young fully expanded leaves of five weeks-old *N. benthamiana* plants. The plants were maintained in a growing chamber at 24°C (light)/20°C (darkness) 16h-light/8h dark photoperiod. Infiltrated leaves were treated either with 0.1-10 mM copper sulfate, 20% ethanol, or water in combination with 0.05% fluvius by spray at three days post infiltration (dpi). The spray was applied to both the adaxial and abaxial surfaces of the leaf.

### Quantification of reporter gene expression

One 8 mm-diameter disc per infiltrated leaf was collected at 4 dpi and stored at –80°C. For analysis, the samples were homogenized, extracted in 180 μl Passive Lysis Buffer (Promega), and centrifugated for 15 min at 14,000 ×*g* and 4°C. An aliquot of 10 μl from the resultant supernatant was transferred to a 96-well plate. As the reporter gene that we used to determine gene expression was the Firefly Luciferase (FLuc), quantification of gene expression was done by measuring the luminescence generated by FLuc. For this measurement, the Dual-Glo^®^ Luciferase Assay System (Promega) was used. The 10 μl of crude extract were mixed with 40 μl of LARII and the FLuc activity was measured by a GloMax 96 Microplate Luminometer (Promega), setting a 2-seconds delay and a 10-seconds measurement. After measuring, 40 μl of Stop&Glo reagent were added per sample and Renilla Luciferase (RLuc) activity was quantified with the same protocol. FLuc/RLuc ratios were calculated as the mean value of the three agroinfiltrated leaves from the same plant. Relative transcriptional activities were expressed as the FLuc/RLuc ratios in each sample and normalized to the FLuc/RLuc ratios of a reference reporter (GB1116).

### Real-time quantitative PCR analysis

Total RNA was isolated from 100 mg of fresh leaf tissue harvested at 5 dpi by using the GeneJET Plant RNA Purification Kit (Thermo Fisher Scientific). The RNA was treated with rDNase (Invitrogen) before cDNA synthesis. The cDNA was synthesized from 1 μg of total RNA using the PrimeScript™ RT-PCR Kit (Takara). Three technical replicates were measured for gene expression levels in each cDNA sample. The SYBR^®^ fluorescent dye (TB Green^®^ Premix Ex Taq™, Takara) was employed for reporting the amplification of cDNA in the Applied biosystem 7500 Fast Real-Time PCR system with specific primer pairs for DFR gene (Supplementary Table 2). F-BOX gene was amplified as an internal reference (Liu *et al*., 2012) for determining DFR gene expression in each sample (ΔCt). The differential DFR gene expression levels were calculated according to the comparative ΔΔCt method as the ratio between the samples agroinfiltrated with the dCasEV2.1 and a gRNA targeting it to the DFR promoter (gDFR) and those including the same construct but an unrelated gRNA (gMTB), both for copper and water treatments (Livak & Schmittgen, 2001).

## LIST OF ABBREVIATIONS

PTA: programmable transcriptional activators
CI-switch: copper inducible-switch (CUP2:Gal4)
CuSO_4_: copper sulfate
dCas9: nuclease-inactivated Cas9 protein
gRNA: guide RNA
dCasEV2.1: dCas9:EDLL + MS2:VPR + gRNA
CI/dCasEV2.1: copper inducible dCasEV2.1
CBS: copper binding site
CUP2: copper responsive factor
TU: transcriptional unit
DFR: NADPH-dependent dihydroflavonol reductase
FLuc: Firefly luciferase
RLuc: Renilla luciferase
tRNA: transfer RNA
RTA: relative transcriptional activity
qRT-PCR: quantitative real-time PCR
Pol-II: RNA polymerase II
AlcR: ethanol-responsive transcriptional factor
Cs: constitutive transcription
Cu: copper-regulated transcription

## DECLARATIONS

### Ethics approval and consent to participate

Not applicable. This article does not contain any studies with human participants or animals performed by any of the authors.

### Consent for publication

Not applicable

### Availability of data and materials

The data and materials generated during the current study are available from the corresponding author on reasonable request.

### Competing interests

The authors declare that they have no competing interests.

### Funding

This work has been funded by EU Horizon 2020 Project Newcotiana Grant 760331 and PID2019-108203RB-100 Plan Nacional I+D, Spanish Ministry of Economy and Competitiveness. Vazquez-Vilar, M. is recipient of APOSTD/2020/096 (Generalitat Valenciana and Fondo Social Europeo post-doctoral grant). García-Pérez, E. is recipient of ACIF-2020 fellowship (Generalitat Valenciana). Diego-Martin, B. and Moreno-Gimenez, E. are recipients of FPU fellowships.

### Author’ contributions

E. G-P., B. D-M, A. Q-R., M. V-V. designed the experiments. E. G-P., B. D-M, A. Q-R., E. M-G. conducted the experiments. E. G-P., M. V-V, D. O. drafted the manuscript. All the authors revised and approved the final manuscript.

## Notes

### Competing Interest Statement

The authors have declared no competing interest.

