## Supplementary tables for "A copper switch for inducing CRISPR/Cas9-based transcriptional activation tightly regulates gene expression in *Nicotiana benthamiana*"

Supplementary Table 1: GB level 1 and >1 transcriptional units and modules. Sequences can be found at http://www.gbcloning.upv.es/ using the GB number or GB name.

| GB number | Construct | GB name | Description |
|---|---|---|---|
| GB0107 | Stuffer fragment | pEGB SF | Used as a stuffer construct to get the same infiltration DO for every *Agrobacterium* mix. |
| GB1116 | pNos:Luciferase:tNos-35S:Renilla:tNos35S:P19:tNos | pEGB PNos:Luciferase:TNos-SF-35S:Renilla:TNos-35S:P19:TNos-SF | Module for the expression of the Firefly Luciferase, the Renilla Luciferase and the P19 silencing suppressor (standard reference in pCambia for luciferase experiments). |
| GB1160 | SlDFR:Luciferase:tNos-35S:Renilla:tNos-35S:P19: tNos | pEGB SlDFR:Luc:TNos-SF-35S:Renilla:TNos-35S:P19:TNos | Module for the expression of the Firefly Luciferase gene driven by the *S. lycopersicum* DFR promoter, the Renilla Luciferase gene and the silencing suppressor P19 driven by the 35s promoter and the Nos terminator. |
| GB1541 | pNos:CUP2:Gal4:tNos | pEGB_3alpha2 PNos:Cup2:Gal4AD:Tnos | Transcriptional unit for constitutive expression of the transcriptional activator protein CUP2 in C-terminal fusion with GAL4 activation domain. Binds to specific DNA operator (CBS) in the presence of copper ions and promotes transcription. |
| GB1838 | U626:gRNA-150 SlDFR: scf2.1 | 3alpha1_U6-26-1gRNA-DFR F6x2 MS2scf | Module for the expression by polymerase III of a guide RNA targeting the DFR promoter with two MS2 aptamer copies in the 3' of the scaffold. |
| GB2070 | U626:multigRNA SlMTB: scf2.1 | pDGB3_Alpha1_U6-26-5gRNA MTB-F6x2_U6-26-4gRNA MTB-F6x2_U6-26-3gRNA MTB-F6x2_U6-26-2gRNA MTB-F6x2 | Module for the expression of 5,4,3,2 gRNA multiplexing targeting the *S. lycopersicum* Mtb promoter with two Ms2 aptamer copies in the 3' of the scaffold (scaffold 2.1). |
| GB2085 | 35S:MS2:VPR:tNos-35S:dCas9:EDLL:tNos | pDGB3_Omega1_35s-Ms2:VPR-Tnos-35s-dCas9:EDLL-Tnos | Module for the constitutive expression of MS2 fused to VPR and dCas9 fused to EDLL. |
| GB2170 | U626:multigRNA NbDFR: scf2.1 | 3Alpha2_U6-26-sgRNANbDFR -85,-198,-268 F6x2 Multiplex | Multiplex construct with F6x2 aptamer in scaffold to target -85,-198,-268 position of *Nicotiana benthamiana* DFR promoter. |
| GB2725 | pNOS:CUP2:VPR:tNos | 3a2:pNOS:CUP2:VPR:tNOS | Transcriptional unit for the expression of CUP2 (copper responsive transcription factor) fused to VPR under the regulation of a NOS promoter and terminator. |
| GB2727 | pNOS:CUP2:TV:tNos | 3a2:pNOS:CUP2:Tv:tNOS | Transcriptional unit for the expression of CUP2 (copper responsive transcription factor) fused to TV under the regulation of a NOS promoter and terminator. |
| GB2740 | CBS4:miniDFR:Luciferase:t35S-35S:Renilla:tNos-35S:P19:tNos | 3o2:CBS4:DFRmin:Luc:t35S-SF-35S:Ren:tNOS-35S:P19:tNOS | Module for the copper inducible expression of luciferase reporter under the regulation of CBS4 operator and the minimal DFR promoter. |
| GB3754 | 35S:MS2:VPR:tNos-CBS4:miniDFR:dCas9:EDLL:tNos | o1_35S:MS2:VPR:Tnos-CBS:miniDFR:dCas9:EDLL:Tnos | Module for simple copper-regulated dCasEV2.1. Copper-regulated dCas9:EDLL with CBS operator + minimal DFR promoter and constitutive MS2:VPR with 35S. |

| | | | |
|---|---|---|---|
| GB3771 | CBS4:mini35S:MS2:VPR:tNos-CBS4:mini35S: dCas9:EDLL:tNos | 3o1:CBS4:35Smin:omegaUTR:MS2:VPR:tNOS-CBS4:35Smin:omegaUTR:dCas9:EDLL:tNOS | Module for double copper-regulated dCasEV2.1. dCas9:EDLL with CBS operator + minimal 35S promoter and MS2:VPR with CBS operator + mini35S promoter. |
| GB3774 | CBS4:miniDFR:MS2:VPR:tNos-CBS4:miniDFR: dCas9:EDLL:tNos | 3o1:CBS4:DFRmin:MS2:VPR:tNOS-CBS4:DFRmin:dCas9:EDLL:tNOS | Module for double copper-regulated dCasEV2.1. dCas9:EDLL with CBS operator + minimal DFR promoter and MS2:VPR with CBS operator + mini35S promoter. |
| GB3786 | CBS4:miniDFR:SlDFR150: scRNA2.1-tRNA:tNOS | 3a1:CBS4:DFRmin:tRNA-SlDFR-150-scRNA2.1-tRNA:tNOS | Transcriptional unit for copper-inducible expression of a sgRNA targeting SlDFR promoter (position -150) under the regulation of the CBS4 operator and DFRmin promoter. This guide RNA contains a scRNA2.1 useful for transcriptional regulation. |
| GB3787 | AlcA:miniDFR:SlDFR150: scRNA2.1-tRNA:tNOS | 3a1:AlcA:DFRmin:SlDFR-150-scRNA2.1-tRNA:tNOS | Transcriptional unit for ethanol-inducible expression of a sgRNA targeting SlDFR promoter (position -150) under the regulation of the AlcA operator and DFRmin promoter. This guide RNA contains a scRNA2.1 useful for transcriptional regulation. |
| GB3788 | pNos:gRNA-150 SlDFR: scf2.1 | 3a1:pNOS:tRNA-SlDFR-150-scRNA2.1-tRNA:tNOS | Transcriptional unit for the expression of a sgRNA targeting SlDFR promoter (position -150) under the regulation of a NOS promoter. This guide RNA contains a scRNA2.1 useful for transcriptional regulation. |
| GB3789 | p35S: gRNA-150 SlDFR: scf2.1 | 3a1:p35S:tRNA:SlDFR -150-scRNA2.1:tNOS | Transcriptional unit for the expression of a sgRNA targeting SlDFR promoter (position -150) under the regulation of a 35S promoter. This guide RNA contains a scRNA2.1 useful for transcriptional regulation. |
| GB3813 | pNOS:CUP2:VP16:tNos | 3a2:pNOS:CUP2:VP16:tNOS | Transcriptional unit for the expression of CUP2 (copper responsive transcription factor) fused to VP16 under the regulation of the NOS promoter and terminator. |
| GB3820 | pNOS:CUP2:Gal4:tNos-pNOS:AlcR:tNos | 3a1_pNOS:CUP2:Gal4:tNOS-pNOS:AlcR:tNOS | Copper and ethanol responsive transcription factors that binds to CBS operator and AlcA operator respectively. |

Supplementary Table 2: Primer pairs for RT-qPCR

| Target gene | Sequence 5'-3' |
|---|---|
| *Nb*DFR | TTCATCTGCGCATCCCATCA |
| | TCCCTACTGAGTTTAAAGGTATCGA |
| *Nb*F-Box | TTGGAAACTCTCTCCCCACTTG |
| | GCTCATTGTTGGATGGGTACCT |
